# EUROPEAN SUBTYPE TICK-BORNE ENCEPHALITIS VIRUS EMERGENCE IN SOUTH KOREA (2005 – 2014) CAN BE ASSOCIATED WITH HUMAN ACTIVITY

**DOI:** 10.1101/2024.11.29.626021

**Authors:** S.Y. Kovalev

## Abstract

Tick-borne encephalitis virus (TBEV) is the etiological agent of tick-borne encephalitis. TBEV is maintained in nature by the ixodid ticks *Ixodes ricinus* and *I. persulcatus*, which are its hosts and main vectors. The classification distinguishes three TBEV subtypes: Far Eastern, European (TBEV-Eu) and Siberian. Each subtype is characterized by its own range, reflected in its name. Nevertheless, numerous findings of TBEV subtypes in non-endemic areas have been reported. This phenomenon may be the result of virus introduction due to natural causes such as bird and mammal migrations. Detection of TBEV-Eu in South Korea in 2005 and in Siberia in the second half of the 20th century are significantly different from other cases of virus introduction. This paper presents the results of a comprehensive study including phylogenetic analysis, clusteron approach, establishing specificity of viral population dynamics, etc. The age of viral populations and the probable time of their emergence in non-endemic regions were determined. Using the example of TBEV-Eu in South Korea, we conclude that there is negative population dynamics of the virus after its introduction. This conclusion, in conjunction with several other features, indicates that human activity played a pivotal role in the TBEV-Eu introduction into South Korea and Siberia.

## Introduction

Tick-borne encephalitis is a vector-borne natural focal infection which causes damage to the central nervous system and is often associated with severe complications in humans, up to and including lethal outcomes. The etiologic agent of this infection is the tick-borne encephalitis virus (TBEV), which belongs to the *Orthoflavivirus* genus of the Flaviviridae family. The TBEV genome is a positive single-stranded RNA molecule, approximately 11,000 bases in length. It encodes three structural proteins (the core [C], pre-membrane [prM], and envelope [E] proteins), and seven non-structural (NS) proteins (NS1, NS2A, NS2B, NS3, NS4A, NS4B, and NS5) within a single open reading frame ^1^.

In accordance with the prevailing taxonomic classification, TBEV is designated as a member of the mammalian tick-borne flavivirus group and the arbovirus ecological group. TBEV is traditionally classified into three subtypes, namely Far Eastern (TBEV-FE), Siberian (TBEV-Sib), and European (TBEV-Eu) ^2^. Recently, two new subtypes were identified, the 886-84-like strain, which is divided from TBEV-FE and named Baikalian (TBEV-Bkl), and Himalayan (TBEV-Him) which has been discovered in *Marmota himalayana* in the Qinghai-Tibet Plateau in China ^3,4^.

TBEV is maintained in nature by ixodid ticks, which are its main hosts, and by vertebrates, through which the virus is transmitted from an infected tick to an uninfected one. TBEV infection has been documented in 18 species of ixodid ticks ^5,6^. However, only two species belonging to the genus Ixodes serve as hosts for TBEV: the taiga tick *Ixodes persulcatus* (Schulze, 1930) and the castor bean tick *I. ricinus* (Linnaeus, 1758) (Acari: Ixodidae). The distribution of TBEV in Eurasia encompasses the ranges of these two species.

TBEV-Eu is associated with *I. ricinus*, whereas all other virus subtypes are associated with *I. persulcatus* . In addition, *I. persulcatus* is a polytypic species and has a pronounced intraspecific structure. Based on morphometric data of all active phases of ontogenesis, two geographically distinct morphotypes are reliably distinguished in it: the ‘Primorsky’ or ‘Eastern’ morphotype and the rest of the range territory - ‘Western’ ^7^. TBEV-FE is associated with the Primorsky morphotype of *I. persulcatus*, while TBEV-Sib is associated with the ‘Western’ morphotype. Thus, the range of TBEV extends in an almost continuous strip across the southern part of the forest zone of extratropical Eurasia from the Atlantic Ocean and the Mediterranean Sea to the Pacific Ocean, including part of Sakhalin Island and the northern part of Hokkaido Island ^5,8^. The main part of the virus range is located within Russia; in addition, it partially or completely covers the territories of 27 European and 7 Asian countries ^9,10^.

The main route of transmission of the virus from an infected tick to an uninfected one is through feeding together on the same animal, although the animal does not necessarily need to develop viremia ^11^. More than 300 different vertebrate species (wild and domestic mammals, birds and reptiles) are feeders for ixodid ticks and therefore take part in the maintenance of natural foci of TBE ^12^. From a general biological standpoint, ticks *I. persulcatus* and *I. ricinus* are hosts for TBEV, while vertebrates (mainly small mammals) act as a host for the tick and are an “intermediary” in the transmission of the virus between ticks. To date, no significant adaptations of TBEV to a specific vertebrate species have been found. The exception is the muskrat (*Ondatra zibethicus*), a semi-aquatic mammal, the adaptation of TBEV to which led to the emergence of the Omsk hemorrhagic fever virus ^13^.

The geographical approach applied to the names of TBEV subtypes correspond to their natural (endemic) ranges. However, some researchers believe that the geographical names of TBEV genotypes do not reflect their strict confinement to a particular region. They correspond only to the higher frequency of occurrence of a particular subtype in a particular region. At least TBEV-Eu and TBEV-FE are found throughout the entire range, but their relative frequency in different parts of the range is not equal ^14^. TBEV-Eu has been found in Altai (Western Siberia, Russia) ^15^, Irkutsk Region (Eastern Siberia, Russia) ^16^, and the Republic of Korea (South Korea) ^17,18^. On the other hand, TBEV-FE was detected in Irkutsk Region (Eastern Siberia) ^16^, Kemerovo and Novosibirsk Regions (Western Siberia) ^19,20^, and Sverdlovsk Region (Middle Urals) ^19,21^. There is another point of view: TBEV subtypes found in non-endemic areas are the result of virus introduction. The introduction of the virus can occur both due to natural causes, such as animal migration (birds and mammals) and/or climatic changes, and due to anthropogenic factors, i.e. because of direct or indirect human involvement.

This paper examines a specific case of TBEV-Eu introduction into South Korea (2005-2014), using data from the introduction of this subtype into Western and Eastern Siberia. Reasoned arguments are given in favor of the decisive role of human activity in the introduction of the virus.

## Materials and methods

Forty TBEV-Eu sequences from GenBank containing a fragment of the E gene with a length of 454 nucleotides were included in the study. All virus strains were isolated from non-endemic regions. South Korea (n=18), Western Siberia: Altai Krai, neighborhood of Zmeinogorsk (n=5), Altai Republic (n=1) and Eastern Siberia: Irkutsk Region (n=16) (Figure 1). All available information on the strains is given in Table S1. Sequences not corresponding to the E gene fragment were not included in the study.

**Figure 1.**
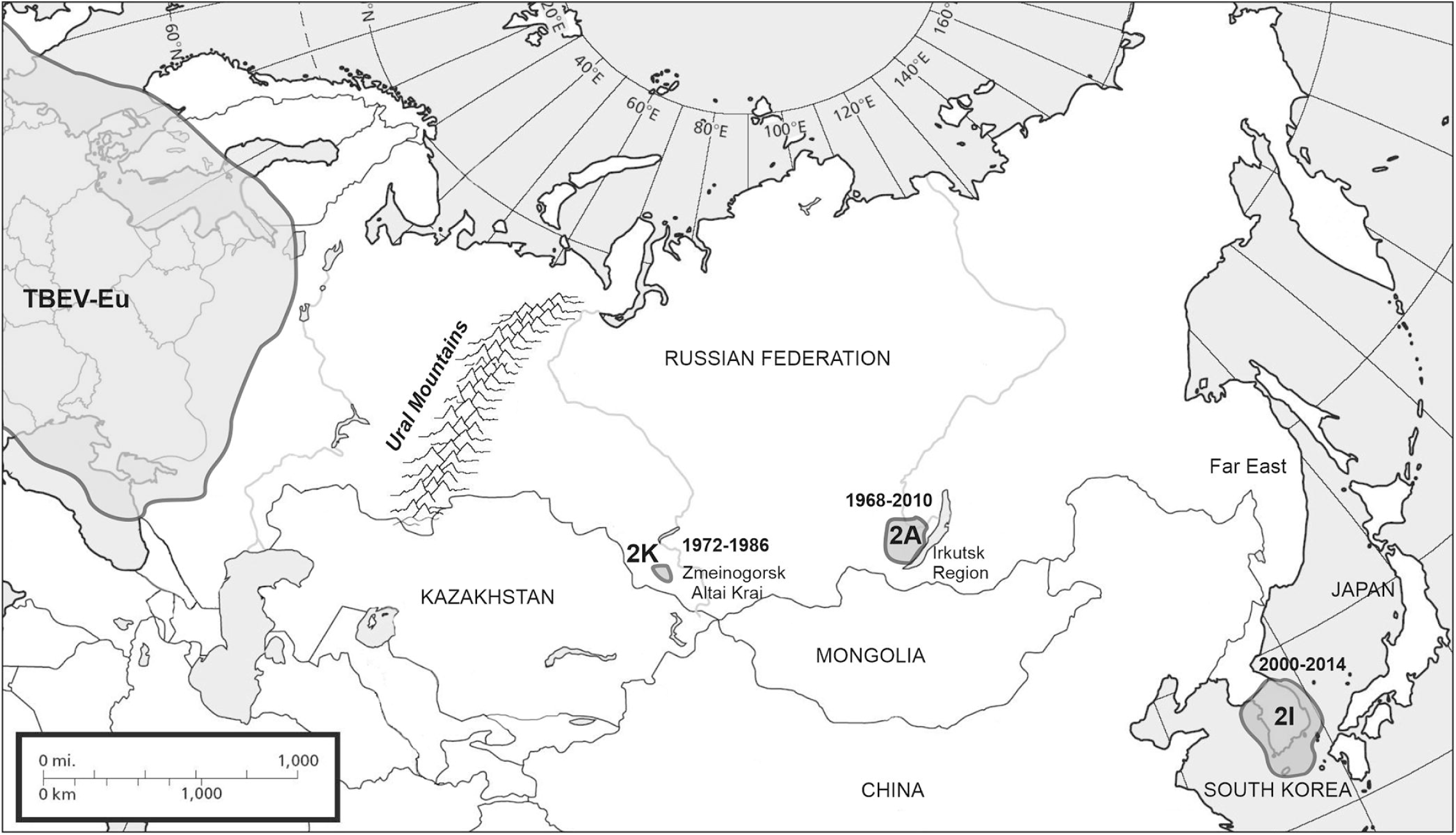
Map of localizations of TBEV-Eu populations in South Korea, Western and Eastern Siberia. The dates indicate the period of their existence (from the beginning of introduction to the isolation of the last strain). 2K, 2A, and 2I are the clusterons to which the strains belong.

Multiple sequence alignment was performed using the MUSCLE software module, part of the MEGA software package, v.11.0.13. Distance tables were generated in MEGA to determine pairwise sequence identity.

Phylogenetic analysis was performed for all TBEV-Eu sequences from GenBank. It included the nucleotide sequences of the E gene fragment (from 309 to 762 nt according to the Zausaev strain GenBank: AF527415) and the deduced amino acid sequences (from 104 to 254 aa) of the corresponding protein. The phylogenetic tree was constructed by the Neighbor-Joining method for TBEV gene E fragment sequences using the p-distance model ^22^ in the MEGA11 software ^23^. The TBEV-FE sequence (strain SofjinKSY_JF819648) was taken as an outgroup. Phylogenetic relationships between different TBEV isolates were confirmed by bootstrap analysis with values >50%. Phylogenetic networks were reconstructed for clusterons using Phylogenetic Network Software v. 10.2 (www.fluxus-engineering.com) with the Median-joining algorithm ^24^. The clusterons to which the sequences belonged were determined by the on-line TBEV Analyzer v.3.0 software (https://tbev.viroinformatics.com/) ^25^. Sequences of the TBEV were grouped into clusterons, sharing the identical amino acid signature of the E protein fragment and being phylogenetically related, according to the approach proposed earlier ^26^.

Evolutionary ages of groups of TBEV-Eu strains included in clusterons were calculated based on the previously determined rate of nucleotide substitution, 1.56 ± 0.29 × 10^−4^ synonymous substitutions per site per year ^27^.

The dynamics of TBEV-Eu infected ticks by year was calculated in Microsoft Excel (Table S2).

## Results

### Phylogenetic and clusteron analysis of TBEV-Eu

Phylogenetic analysis of the E gene fragment of TBEV-Eu strains from South Korea showed that they are part of the same clade (Figure 2A). Eleven of the 18 TBEV-Eu strains had an identical amino acid sequence of the protein E fragment and were part of clusteron 2I in the TBEV-Eu clusteron structure (Figure 2B). The derived clusteron 2I differs from the clusteron-founder 2A by a single amino acid substitution of E122G (Figure 2A).

**Figure 2.**
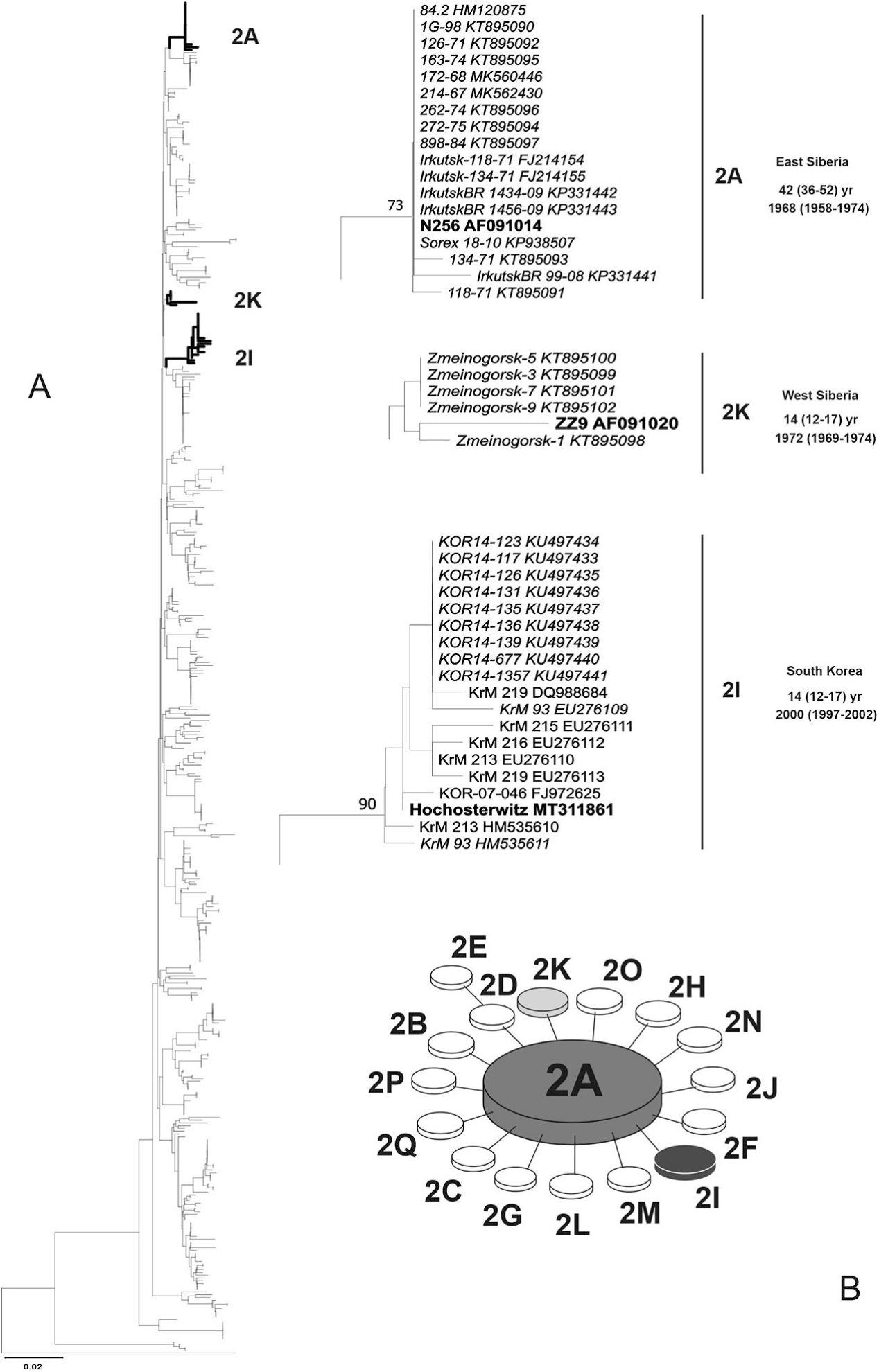
Phylogenetic analysis of all TBEV-Eu strains. A. Dendrogram based on the nucleotide sequences of the E gene fragment. Strains found in South Korea and Siberia are shown to scale with the age of the viral population and the estimated time of introduction. Strains belonging to clusterons 2A, 2K, and 2I are shown in italics. Prototypic strains are shown in bold. B. Clusteron structure of TBEV-Eu.

Sixteen TBEV-Eu sequences from Eastern Siberia are also phylogenetically closely related. In the TBEV-Eu clusteron structure, they are part of the clusteron-founder 2A (Figure 2B).

Five of the six nucleotide sequences of the virus isolated in Western Siberia are also phylogenetically closely related. All have an identical amino acid sequence of protein fragment E and form a derivative clusteron 2K. Clusteron 2K differs from the clusteron-founder 2A by a single amino acid substitution of A146V (Figure 2). The exception is strain 84.2 (HM120875), which is part of clusteron 2A and is phylogenetically closely related to TBEV-Eu from Eastern Siberia (Figure 2, Table S1)

### Dynamics of virus-infected ticks over the observation period

The first evidence of the presence of TBEV-Eu in South Korea was obtained in 2005 ^17^. Over the next 10 years, regular observations were carried out, which made it possible to assess the dynamics of virus-infected ticks (Figure 3). The initial infection rate was 0.63%. However, in 2014, there was a marked decline, with a reduction of over 10-fold to 0.043% (Table S2) ^28–31^. The dynamics during observations fits into a power function with a trend y = 0.6606x-1.153 (R² = 0.9711) (Figure 3). It is not possible to assess the dynamics of virus-infected ticks for Western and Eastern Siberia due to the lack of data necessary for calculations.

**Figure 3.**
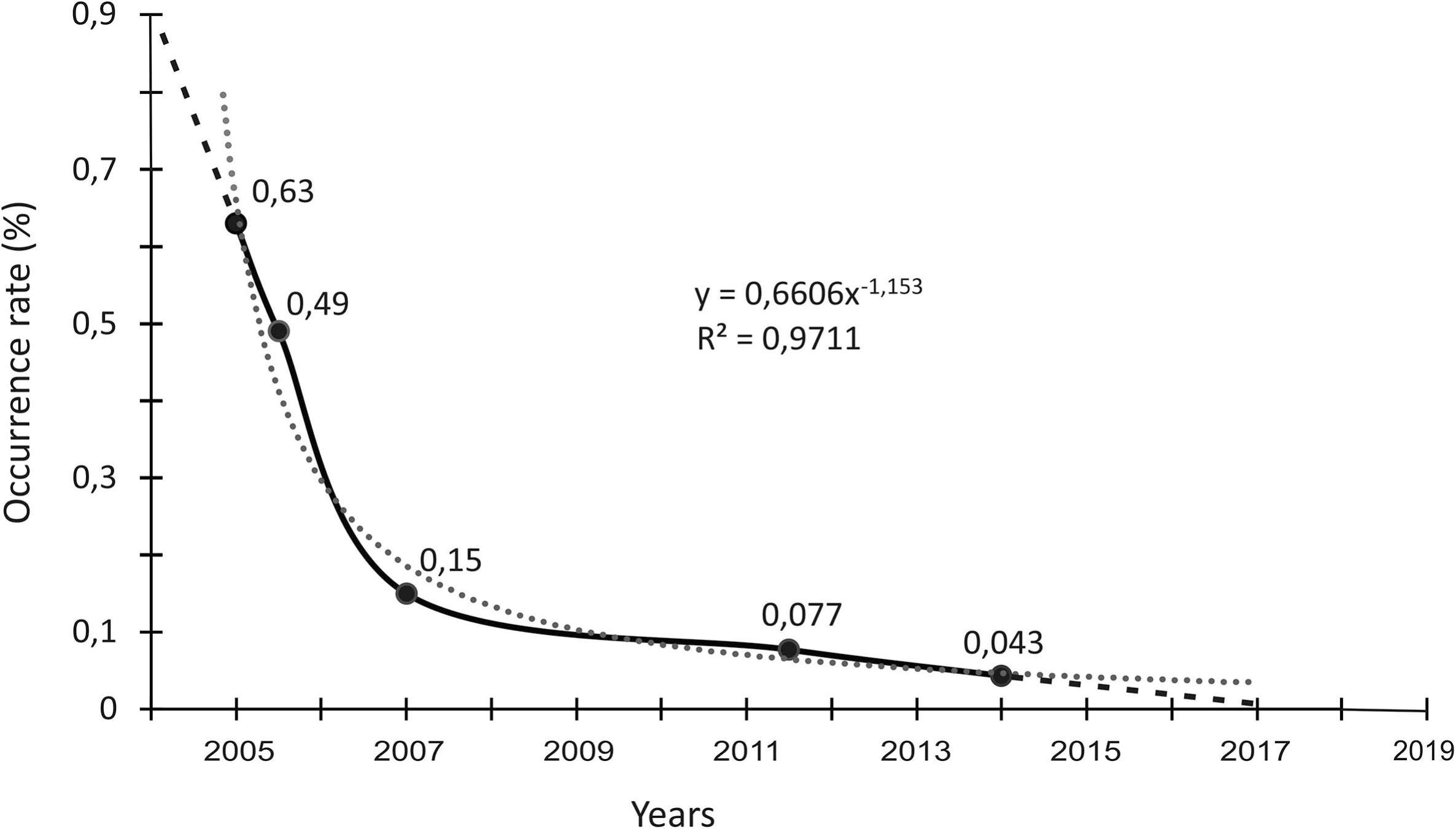
Dynamics of ticks infected of TBEV-Eu in South Korea for the period 2005–2014. The solid line is the ratio of the number of positive TBEV samples to the total number of ticks taken in the study. The dotted line is the estimated trend of this dynamic. R^2^ – approximation reliability value.

### Age estimation of non-endemic TBEV-Eu populations

The calculated age of the Korean TBEV-Eu population, based on the genetic distance between eleven strains of clusteron 2I and the rate of nucleotide substitutions per site per year, was 14 (12–17) years. The age of the West Siberian TBEV-Eu population calculated for clusteron 2K was 14 (12–17) years, and in the East Siberian (2A) population was 42 (36–52) years.

## Discussion

The discovery of TBEV-Eu in South Korea in 2005 was a sensation because it broke the existing assumptions about the confinement of TBEV subtypes to specific geographical regions in Europe, Siberia and the Far East. TBEV subtypes have been detected in non-endemic areas previously, but they were few and localized. They could be explained with some reservations by natural causes. For example, the introduction of infected ticks by migratory birds or mammalian migrations. However, the detection of TBEV-Eu in South Korea led some researchers to think that TBEV subtypes naturally exist in non-endemic regions, and thus they extended the range of individual subtypes to the entire range of the virus ^32^. The results of long-term observations of TBEV-Eu in South Korea allowed us to analyze this phenomenon and come to important conclusions.

### TBEV-Eu emerged concurrently in multiple locations across South Korea

Based on the smoothed shape of the TBEV-Eu occurrence dynamics curve (Figure 3), it can be concluded that the introduction was a single event, or it would have contained fluctuations associated with repeated introductions.

The dynamics of virus-infected ticks allow for the accurate establishment of the time of the virus introduction (Figure 3). The prevalence of TBEV-Eu in *I. ricinus* in Europe is estimated to be between 0.02% and 2.0% ^33^. If we assume that at the outset of the introduction, the prevalence of the virus in ticks was comparable to the midpoint value observed in Europe, then the introduction of the virus into South Korea could have occurred at the very beginning of the 2000s, no later than 2003. An alternative approach, based on the calculation of clusteron 2I age, indicated that the introduction of TBEV-Eu occurred in 2000 (1997-2002). The concordance in time estimation between the two independent methods provides a reliable date for the virus introduction into South Korea, which can be placed between 2000 and 2003.

The scale of the TBEV-Eu introduction is unprecedented. The virus has been identified in seven of the nine provinces of South Korea, encompassing nearly the entire country’s territory of approximately 100,000 square kilometers ^34^. It is noteworthy that the virus has been identified on Jeju Island (Jeju-do Province), situated in the Korea Strait over 80 kilometers from the mainland (Figure 4) ^35^. Despite the extensive geographical scope, the introduction of the virus was confined to the territory of a single country. There have been no documented instances of the virus being detected in neighboring countries, including China, Japan and Russia (Primorsky Krai), where only the TBEV-FE is found.

**Figure 4.**
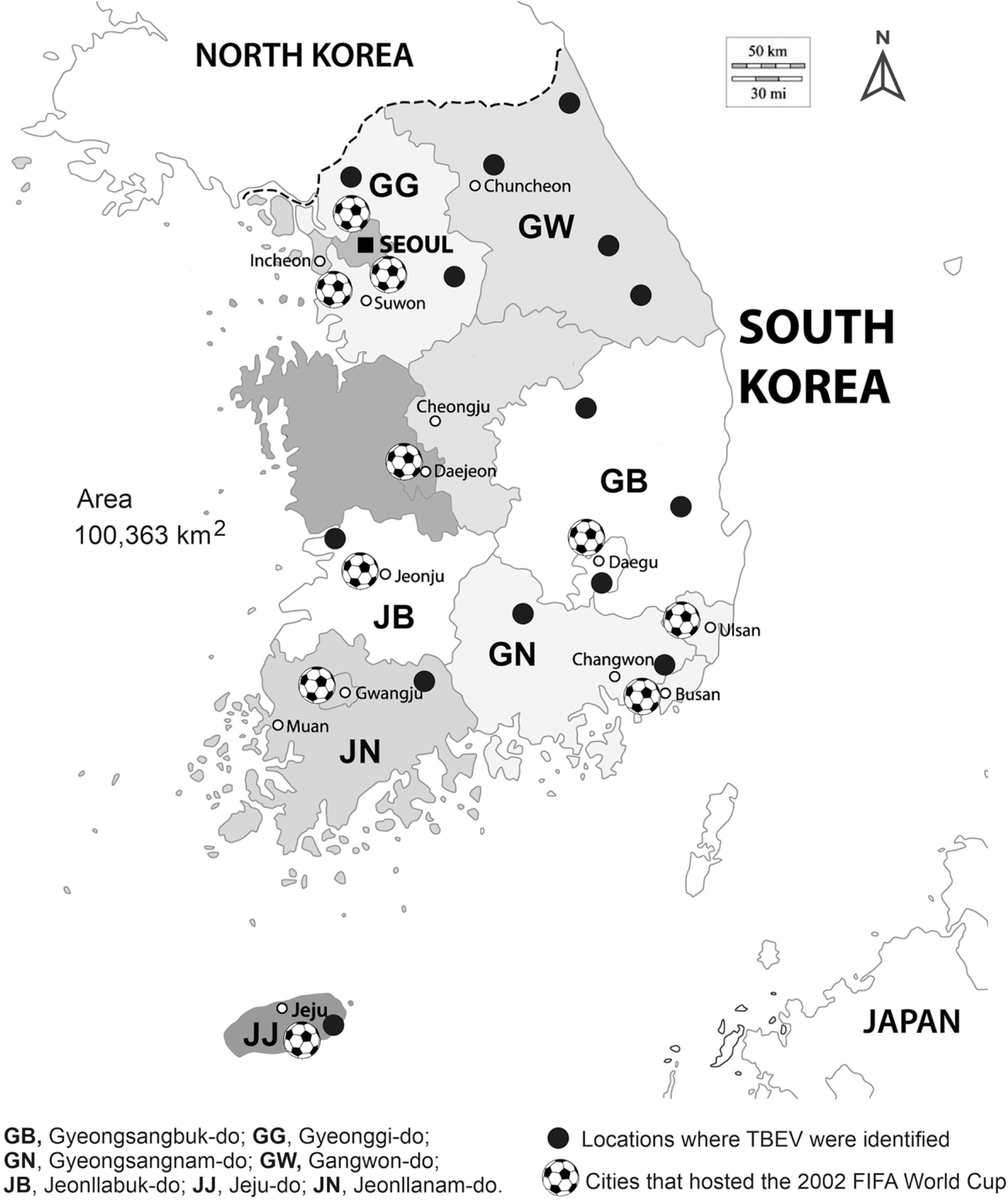
Map of South Korea showing the locations of TBEV-Eu detections and the cities where the 2002 FIFA World Cup matches were held.

The introduction of TBEV-Eu into Eastern and Western Siberia, calculated from the age of the virus clusterons 2A and 2K, is estimated to have occurred around 1968 (1958-1974) and 1972 (12-17), respectively. In a previously published paper, the time of TBEV-Eu introduction into Eastern Siberia was dated to approximately 1928 (1900-1950) ^36^. Despite the discrepancies in determining the time of introduction, it can be stated that a recent introduction of TBEV-Eu into Siberia has occurred.

### TBEV-Eu in South Korea has now disappeared

During the ten years of observation (2005-2014), the occurrence of the virus in South Korea dropped more than 10-fold. The dynamics of ticks infected of TBEV-Eu shown in Figure 3 suggest the disappearance of TBEV-Eu in South Korea by the late twenties of this century. This conclusion is corroborated by the absence of any detection of the virus in the subsequent ten-year period (2015-2024). The reason for the virus disappearance appears to be the absence of the natural arthropod host tick *I. ricinus* to which the virus is adapted. TBEV-Eu in South Korea has been isolated from the dominant tick species of the genus *Haemaphysalis* (*H. longicornis, H. flava, H. japonica*) and one species of the genus *Ixodes* (*I. nipponensis*), which accounts for about 1%. At the same time, the specific occurrence of the virus in *I. nipponensis* is more than 10 times higher than in *Haemaphysalis spp* . ^28–31^. This indicates that the virus was maintained mainly by *I. nipponensis*, which phylogenetically belongs to the Palearctic group *I. ricinus - I. persulcatus* ^37,38^. The low abundance of *I. nipponensis*, and its inability to maintain the virus as effectively in nature as the natural host, *I. ricinus*, led to the rapid disappearance of TBEV-Eu in South Korea.

The shape of curve shown in Figure 3 leads to the conceptual conclusion that any introduction of TBEV subtypes into non-endemic areas will be accompanied by negative dynamics of the virus population. In other words, such an introduction will inevitably lead to the disappearance of the introduced virus. The duration of the existence of the introduced subtype of the virus depends mainly on two parameters. Firstly, the evolutionary relatedness of ticks belonging to the Ixodidae family in a non-endemic area to the natural host tick of the virus, as well as the proportion of these ticks among other ixodid ticks, must be considered. Secondly, the scale of introduction, defined as the number of viruses introduced, must be considered. TBEV-Eu existed in Eastern Siberia for about 42 years and was no longer detected after 2010 ^39^ (Figure 1). The three-fold duration of TBEV-Eu presence in Eastern Siberia compared to South Korea can be explained by the dominance of *I. persulcatus* ticks ^40^. This tick is a close relative of the natural host of TBEV-Eu, the tick *I. ricinus* .

### TBEV-Eu strains from South Korea, as well as from Western and Eastern Siberia, originate from laboratory strains and have low genetic variability

At first glance, there is nothing in common between the TBEV-Eu strains from South Korea, Western Siberia and Eastern Siberia. They belong to different clades (Figure 2A), belong to different clusterons in the clusteron structure (Figure 2B), and differ in the place and time of introduction (Figure 1). The only thing they have in common is insignificant genetic variability (Figure 2), as well as close affinity with the ‘old’ laboratory strains from which they derive. The only European strain related to the South Korean strain is the Hochosterwitz strain (MT311861) isolated in Austria in 1971 from the tick *I. ricinus* (Figure 2A, Table S1). The relatedness of the South Korean strains to the Hochosterwitz strain (MT311861) was also previously shown by full genome sequence analysis ^41^. The peculiarity of this strain is its uniqueness, i.e. the absence of phylogenetically similar strains in Europe. It should be noted that the Hochosterwitz strain (MT311861) is a laboratory strain ^42^.

The TBEV-Eu strains isolated in Western and Eastern Siberia were also found to be closely related to laboratory strains Absettarov (AF091005) and N256 (AF091014), respectively (Figure 2, Table S1). This phylogenetic relatedness was additionally confirmed by full genome sequence analysis ^15^. Therefore, there is a general tendency for TBEV-Eu strains from Asia to be associated with laboratory strains.

### The role of the human activity in the introduction of TBEV-Eu in South Korea and Siberia

The introduction of TBEV-Eu in South Korea and Siberia has been attributed to natural causes, namely the transfer of virus-infected ticks by migratory birds or small mammals. This hypothesis is unlikely to be correct for several reasons. Firstly, the introduction of the virus is not widespread, occurring instead as isolated cases. It is practically difficult to detect because of the rapid decline in virus abundance in the absence of a natural arthropod host (Figure 3). Secondly, the introduction is not selective and is random in nature. In all three cases considered, the introduced virus originated from ’old’ laboratory strains of TBEV-Eu. The probability that South Korea, and especially Jeju Island, will encounter a virus originating from the unique Hochosterwitz strain (MT311861) is negligible. Third, long-distance migratory birds make their seasonal migrations from south to north and back. Birds nesting in the north of Europe do not follow migration routes over Siberia and the Far East and therefore cannot be the cause of TBEV-Eu introduction. On the other hand, short-distance or altitudinal migratory birds cannot cover distances of more than 7,000 km. Interestingly, TBEV has never been detected in South Korea during the entire observation period up to 2005. At the same time, TBEV-FE is widespread in neighboring countries. The probability of introduction of TBEV-FE by migrating birds to South Korea from neighboring countries is several orders of magnitude higher than the introduction of TBEV-Eu from the European continent.

An alternative explanation for the introduction of the virus could be human activity, i.e. direct or indirect human involvement in the transportation of the TBEV-infected ticks into non-endemic areas. The anthropogenic factor has a strong potential to transfer tick-borne flaviviruses over long distances. It is assumed that the introduction of louping ill virus (this virus is phylogenetically close to TBEV-Eu) to Norway was associated with the importation of sheep from the British Isles ^43^. The expansion of new territories of Siberia and the Far East in the XVII-XVIII centuries, the construction of long trunk communication routes, as well as the state Programme for Acclimatization of Game Animals and Birds in the former Soviet Union, had a direct or indirect impact on the introduction of the virus to new territories ^4,21,27^.

It is not known exactly how and under what circumstances the virus was introduced into South Korea, a problem that can only be solved by the Korea Disease Control and Prevention Agency (KDCA). Nevertheless, the introduction of the virus must have occurred because of a specific event that fulfils the following criteria. Firstly, it is necessary to ascertain that the event in question occurred between the years 2000 and 2003. Secondly, the phenomenon had to be of a considerable magnitude, encompassing the entirety of the country, including the island province of Jeju-do. Thirdly, the event in question had to be of an international nature. A single extraordinary event in the history of South Korea aligns with these criteria: the FIFA World Cup, which was held from 31 May to 30 June 2002. It is evident that the sporting event itself cannot be identified as the direct cause of the virus’ introduction. Nevertheless, a potential link may exist between the virus introduction and preparations for football matches, as well as activities providing leisure, recreational, and transport services for tens of thousands of fans and tourists across South Korea. This is corroborated by the fact that TBEV-Eu strains were isolated in six of the seven provinces where football matches were held, including the island province of Jeju-do (Figure 4). It should be noted that this introduction exhibited a national character, as the World Cup matches were held in two countries: South Korea and Japan, yet TBEV-Eu was not detected in Japan.

It is fortunate that no cases of tick-borne encephalitis have been reported in South Korea because of the introduction of TBEV-Eu. However, antibodies to TBEV have been detected in forest-related people ^44,45^. The introduction of TBEV-Eu into Western and Eastern Siberia is probably also associated with human activity, but how exactly it was realized is still difficult to establish due to the lack of required information.

## Conclusion

The natural introduction of TBEV subtypes into non-endemic areas is of little concern, given that it is spontaneous and isolated. However, the introduction of the virus with human involvement may be large-scale and pose a serious threat to public health. TBEV-Eu in South Korea is an example of this. Identifying the causes and ways to prevent such an introduction is a challenge for TBE epidemiologists and others.

## Supporting information

List of nucleotide sequences of TBEV-Eu strains isolated in South Korea and Siberia

Relative abundance of TBEV-Eu positive samples from ticks in different years in South Korea.

## Acknowledgements

The authors are indebted to Dr. Keith Chamberlain (United Kingdom) and Dr. Irina Belyaeva (United Kingdom) for their linguistic help in preparing the article.

