## Supplementary material for "EUROPEAN SUBTYPE TICK-BORNE ENCEPHALITIS VIRUS EMERGENCE IN SOUTH KOREA (2005 – 2014) CAN BE ASSOCIATED WITH HUMAN ACTIVITY": List of nucleotide sequences of TBEV-Eu strains isolated in South Korea and Siberia

**Supplement Table 1.** List of nucleotide sequences of TBEV-Eu strains isolated in South Korea and Siberia

| **#** | **Strain** | **Clusteron** | **GenBank**  **Accession no.** | **Year of isolation** | **Geographical origin** | **Source of isolate** | **References** |
| --- | --- | --- | --- | --- | --- | --- | --- |
| 1. | Zmeinogorsk-1 | 2K | KT895098 | 1986 | Russia: Western Siberia, Altai Krai | *I. persulcatus* | Tkachev et al. (2015) |
| 2. | Zmeinogorsk-3 | 2K | KT895099 | 1986 | Russia: Western Siberia, Altai Krai | *I. persulcatus* | Tkachev et al. (2015) |
| 3. | Zmeinogorsk-5 | 2K | KT895100 | 1986 | Russia: Western Siberia, Altai Krai | *I. persulcatus* | Tkachev et al. (2015) |
| 4. | Zmeinogorsk-7 | 2K | KT895101 | 1986 | Russia: Western Siberia, Altai Krai | *I. persulcatus* | Tkachev et al. (2015) |
| 5. | Zmeinogorsk-9 | 2K | KT895102 | 1986 | Russia: Western Siberia, Altai Krai | *I. persulcatus* | Tkachev et al. (2015) |
| 6. | 84.2 | 2A | HM120875 | 2007 | Russia: Western Siberia, Altai Republic | *I. persulcatus* | Tikhomirov et al. (2010) |
| 7. | Altay-103 | 2* | JQ687275 | 2007 | Russia: Western Siberia, Altai Republic | *I. persulcatus* | Karan et al. (2012) |
| 8. | IrkutskBR_99-08 | 2A | KP331441 | 2008 | Russia: Eastern Siberia, Irkutsk Reg. | *I. persulcatus* | Adelshin et al. (2014) |
| 9. | IrkutskBR_1434-09 | 2A | KP331442 | 2009 | Russia: Eastern Siberia, Irkutsk Reg. | *I. persulcatus* | Adelshin et al. (2014) |
| 10. | IrkutskBR_1456-09 | 2A | KP331443 | 2009 | Russia: Eastern Siberia, Irkutsk Reg. | *I. persulcatus* | Adelshin et al. (2014) |
| 11. | Sorex 18-10 | 2A | KP938507 | 2010 | Russia: Eastern Siberia, Irkutsk Reg | *Sorex sp.* | Adelshin et al. (2015) |
| 12. | 118-71 | 2A | KT895091 | 1971 | Russia: Eastern Siberia, Irkutsk Reg. | *Spermophilus undulatus* | Tkachev et al. (2015) |
| 13. | 126-71 | 2A | KT895092 | 1971 | Russia: Eastern Siberia, Irkutsk Reg. | *I. persulcatus* | Tkachev et al. (2015) |
| 14. | 134-71 | 2A | KT895093 | 1971 | Russia: Eastern Siberia, Irkutsk Reg. | *Spermophilus undulatus* | Tkachev et al. (2015) |
| 15. | 272-75 | 2A | KT895094 | 1975 | Russia: Eastern Siberia, Irkutsk Reg. | *Microtus gregalis* | Tkachev et al. (2015) |
| 16. | 163-74 | 2A | KT895095 | 1974 | Russia: Eastern Siberia, Irkutsk Reg. | *I. persulcatus* | Tkachev et al. (2015) |
| 17. | 262-74 | 2A | KT895096 | 1974 | Russia: Eastern Siberia, Irkutsk Reg. | *I. persulcatus* | Tkachev et al. (2015) |
| 18. | 898-84 | 2A | KT895097 | 1984 | Russia: Eastern Siberia, Irkutsk Reg. | *Myodes rutilus* | Tkachev et al. (2015) |
| 19. | 1G-98 | 2A | KT895090 | 1998 | Russia: Eastern Siberia, Irkutsk Reg. | *Human blood* | Tkachev et al. (2015) |
| 20. | Irkutsk-118-71 | 2A | FJ214154 | 1971 | Russia: Eastern Siberia, Irkutsk Reg. | *Citellus undulatus* | Karan et al. (2008) |
| 21. | Irkutsk-134-71 | 2A | FJ214155 | 1971 | Russia: Eastern Siberia, Irkutsk Reg. | *Citellus undulatus* | Karan et al. (2008) |
| 22. | 172-68 | 2A | MK560446 | 1968 | Russia: Eastern Siberia, Irkutsk Reg. | *Homo sapiens* | Adelshin et al. (2019) |
| 23. | 214-67 | 2A | MK562430 | 1967 | Russia: Eastern Siberia, Irkutsk Reg. | *Homo sapiens* | Lopatovskaya et al. (2019) |
| 24. | KrM 93 | 2I | EU276109 | 2007 | South Korea | *Tick* | Yun et.al. (2007) |
| 25. | KrM 93 | 2I | HM535611 | 2006 | South Korea | *Apodemus agrarius* | Yun et al. (2010) |
| 26. | KOR14-117 | 2I | KU497433 | 2014 | South Korea | *I. nipponensis* | Yun et al. (2016) |
| 27. | KOR14-123 | 2I | KU497434 | 2014 | South Korea | *I. nipponensis* | Yun et al. (2016) |
| 28. | KOR14-126 | 2I | KU497435 | 2014 | South Korea | *Haemaphysalis flava* | Yun et al. (2016) |
| 29. | KOR14-131 | 2I | KU497436 | 2014 | South Korea | *Haemaphysalis longicornis* | Yun et al. (2016) |
| 30. | KOR14-135 | 2I | KU497437 | 2014 | South Korea | *Haemaphysalis flava* | Yun et al. (2016) |
| 31 | KOR14-136 | 2I | KU497438 | 2014 | South Korea | *Haemaphysalis longicornis* | Yun et al. (2016) |
| 32. | KOR14-139 | 2I | KU497439 | 2014 | South Korea | *Haemaphysalis longicornis* | Yun et al. (2016) |
| 33. | KOR14-677 | 2I | KU497440 | 2014 | South Korea | *Haemaphysalis flava* | Yun et al. (2016) |
| 34. | KOR14-1357 | 2I | KU497441 | 2014 | South Korea | *Haemaphysalis longicornis* | Yun et al. (2016) |
| 35. | KOR-07-046 | 2Unique | FJ972625 | 2007 | South Korea: Jeju Island | *Haemaphysalis longicornis* | Chae (2009) |
| 36. | KrM 213 | 2Unique | EU276110 | 2007 | South Korea | *Tick* | Yun et.al. (2007) |
| 37. | KrM 215 | 2Unique | EU276111 | 2007 | South Korea | *Tick* | Yun et.al. (2007) |
| 38. | KrM 216 | 2Unique | EU276112 | 2005 | South Korea | *Tick* | Kim et al. (2006) |
| 39. | KrM 219 | 2Unique | EU276113 | 2005 | South Korea | *Tick* | Kim et al. (2007) |
| 40. | KrM 213 | 2Unique | HM535610 | 2006 | South Korea | *Apodemus agrarius* | Yun et al. (2010) |
| 41. | KrM 219 | 2Unique | DQ988684 | 2006 | South Korea | *Apodemus agrarius* | Kim et al. (2006) |
|  | **Prototype (laboratory) strain** | |  |  |  |  |  |
| 1. | N256 | 2A | AF091014 | 1940 | Belarus: Minsk | *I. ricinus* | Ecker et al. (1999) |
| 2. | Hochosterwitz | 2A | MT311861 | 1971 | Austria: Hochosterwitz | *I. ricinus* | Paulsen et al.(2020) |
| 4. | Absettarov | 2A | AF091005 | 1951 | Russia: Leningrad Region | *Human blood* | Ecker et al. (1999) |
| 5. | Absettarov | 2A | KU885457 | 1951 | Russia: Leningrad Region | *Human blood* | Romanova et al. (2016) |

* Sequences with degenerate nucleotide positions of the E gene fragment were not included in the study.
