## Supplementary material for "EUROPEAN SUBTYPE TICK-BORNE ENCEPHALITIS VIRUS EMERGENCE IN SOUTH KOREA (2005 – 2014) CAN BE ASSOCIATED WITH HUMAN ACTIVITY": Relative abundance of TBEV-Eu positive samples from ticks in different years in South Korea.

**Supplement Table 2.** Relative abundance of TBEV-Eu positive samples from ticks in different years in South Korea.

| **Year of isolation** | **Number of ticks** | **Number of pools** | ***TBEV qPCR*** | **Ratio of** **positive samples to the number of ticks/pools (%)** | **References** |
| --- | --- | --- | --- | --- | --- |
| 2005 | 635 | 38 | 4 | 0.63/10.5 | (Kim et al., 2008) |
| 2005-2006 | 2460 | 197 | 12 | 0.49/6.1 | (Kim et al., 2009) |
| 2007 | 4077 | 649 | 6 | 0.15/0.92 | (Ko et al., 2010) |
| 2011-2012 | 13053 | 1292 | 10 | 0.077/0.78 | (Yun et al., 2012) |
| 2014 | 21158 | 1518 | 9 | 0.043/0.59 | (Yun et al., 2016) |
